# Using FINDeM for rapid, CRISPR-based detection of the emerging salamandrid fungal pathogen, *Batrachochytrium salamandrivorans*

**DOI:** 10.1101/2024.01.08.573879

**Authors:** Brandon D. Hoenig, Philipp Böning, Amadeus Plewnia, Corinne L. Richards-Zawacki

**Author notes:** Corresponding author: Brandon D. Hoenig 4200 Fifth Ave, Pittsburgh, Pennsylvania, USA 15260 7245615563.

## Abstract

The fungal pathogen *Batrachochytrium salamandrivorans* (*Bsal*) is one of two species (the other, *B. dendrobatidis/Bd*) that cause amphibian chytridiomycosis, an emerging infectious disease that has been indicated in the declines of hundreds of amphibian species worldwide. While *Bd* has been near-globally distributed for well over a century, *Bsal* is a more recently emerged pathogen, having been identified just over a decade ago with current impacts localized to salamandrids in parts of Europe. However, because there is concern that *Bsal* will cause widespread declines if introduced to naïve regions – such as the Americas where the greatest diversity of salamandrids exist – it is imperative that widespread testing and monitoring strategies be implemented to mitigate the spread of *Bsal*. As standard diagnostic approaches tend to be expensive, time-consuming, or require specialized instrumentation and training, we have developed a simplified, rapid, CRISPR-based approach for *Bsal*-DNA identification and provide suggestions for its future application.

## Introduction

Emerging Infectious Diseases (EIDs) have been identified as a rising threat to global biodiversity (1), and while they are commonly associated with public health, agriculture, and livestock, EIDs that are specific to wildlife have caused detrimental and irreversible impacts on various plant (2) and animal populations worldwide (3). However, as the direct effects of EIDs on wildlife – as well as the potential indirect effects that may be incurred by humanity (e.g., hunting-related economic impacts of chronic wasting disease on cervids; 4) – are not always apparent, the importance of these wildlife EIDs tend to receive less attention than those that directly impact humans. For amphibians, the most threatened vertebrate group on the planet, wildlife EIDs are among the main factors causing global population declines (5), with two species of fungus – *Batrachochytrium dendrobatidis* (*Bd*) and *B. salamandrivorans* (*Bsal*), the causative agents of amphibian chytridiomycosis (6) – playing a prominent role in these declines (7). While *Bd* is considered the world’s most impactful invasive species (8) and threatens all three orders of amphibians globally (7), *Bsal* has emerged more recently in Europe, where it has posed a threat to the west Palearctic caudate diversity for at least two decades (9, 10). Outside its hypothesized native range in Asia (11), *Bsal* has been observed in four European countries (Belgium, Germany, Spain, the Netherlands) and, as spill-over events to naïve regions have been noticed repeatedly in Europe (12–14), there is great concern that the pathogen may spread to the Americas and threaten salamander species in the taxon’s global biodiversity hotspot (15).

Mitigating the impacts and spread of wildlife EIDs to naïve regions requires timely detection and continued monitoring (16). However, detection of *Bsal* currently relies on direct diagnosis via epidermal histology or the indirect, DNA-based detection of *Bsal* via endpoint or quantitative PCR (9, 17, 18). Unfortunately, each of these methods may take days to weeks to return results and require specialized equipment within designated facilities as well as researchers who have received specific training in these approaches (19); drawbacks that have allowed only limited active pathogen surveillance thus far (20). However, a recently-developed CRISPR-based, molecular diagnostic tool – FINDeM (<u>F</u>ield-deployable, Isothermal, <u>N</u>ucleotide-based <u>De</u>tection <u>M</u>ethod) – has been used to detect *Bd in situ* immediately upon sample collection without requiring specialized lab equipment, extensive training, or vast financial resources (19). Using a rapid DNA extraction method (21) alongside two body-heat inducible reactions – recombinase polymerase amplification to increase concentrations of target DNA (RPA; 22) and a fluorescence-based, CRISPR-Cas12a reporter assay to identify target DNA (23) – FINDeM has been shown to detect single-digit copies of *Bd* DNA in under one hour (19).

Adapting this assay for the detection of *Bsal* would not only allow a more diverse community of conservation researchers and citizen scientists to increase sampling effort in time and range but may also create avenues for more rapid and comprehensive testing to strengthen the clean trade of amphibians (24, 25). Therefore, in this study we demonstrate the use of FINDeM for *Bsal* detection and show its strengths and limitations based on laboratory standards and field-collected samples in direct comparison with qPCR.

## Methods

### *Bsal* FINDeM design and protocol

RPA reaction master mixes used in FINDeM followed (19), deviating only in using a 10uM-each mixture of standard, *Bsal*-targeting qPCR primers – SterF (5’-TGCTCCATCTCCCCCTCTTCA-3’) and SterR (5’-TGAACGCACATTGCACTCTAC-3’) which target a180bp segment of the 5.8S rRNA gene (9). Moreover, in qPCR, a fluorescently-labeled probe is used (SterC; 5’-ACAAGAAAATACTATTGATTCTCAAACAGGCA-3′; 18) to indicate the presence of *Bsal* DNA. FINDeM instead utilizes a guide RNA (gRNA) that is complementary to a 25bp segment of this same probe region, though on the opposite DNA strand (5’-<u>TTTG</u>AGAATCAATAGTATTTTCTTG-3’; protospacer adjacent motif (PAM) underlined).

Subsequently, 4.5uL of the RPA master mix was added to 5uL of each sample and 0.5uL of 280mM magnesium acetate (MgOAc) was added to the sidewall of the reaction tube. Reaction tubes were pulse spun to initiate the amplification reaction and the tubes were incubated at 37°C for 30 minutes. FINDeM DNA detection mixtures followed (19) but instead contained 0.5uL of 1.25uM *Bsal* guide RNA (gRNA; 5’-UAAUUUCUACUAAGUGUAGAUAGAAUCAAUAGUAUUUUCUUG-3’; IDT) per reaction. This mixture (hereafter, ‘CRISPR Mix’) was incubated at ambient temperature for 30 minutes to allow the *Bsal* gRNA and LbCas12a to form a DNA detection complex. After each reaction’s 30-minute incubation period, 2.5uL of the CRISPR Mix was added to the entire 10uL product of the RPA reaction and the final 12.5uL volume incubated at 37°C on a QuantStudio 3 or StepOne Plus (Applied Biosystems™; Waltham, MA, USA) with ROX fluorescence monitored every 30 seconds for 30 minutes. Immediately after this final incubation, 10uL of the final product was moved to a new tube and photographed under 360-nm ultraviolet light with a cellphone camera.

### FINDeM validation with plasmid and zoospore DNA standards

To test the efficacy of the *Bsal* FINDeM assay and assess its limit of detection (LOD), we performed qPCR and FINDeM in duplicate on serially diluted zoospore-extract standards (range: 0.1 – 10,000 Zoospore Genomic Equivalents or ZGEs; water as negative control) and plasmid DNA standards (range: 2.6 – 2,600,000 *Bsal* DNA copies; Qiagen Buffer AE as negative control). For the FINDeM samples, we followed the protocol described above. The qPCR reactions were performed in duplicate following thermal cycling conditions in (26) on a QuantStudio 3 (Applied Biosystems™; Waltham, MA, USA) and prepared in 25uL volumes with 5uL of each *Bsal* standard serving as template, 12.5uL of 2x Sensi-fast Lo-ROX master mix, 300nM of each primer, 100nM of ABY-labeled SterC probe, 400ng/uL of BSA, and a VIC labeled Exo-IPC mix with DNA loaded at 1/3x concentration.

### FINDeM validation with field-collected samples

We also evaluated the efficacy of qPCR and FINDeM in duplicate on paired swab samples from 22 newts of three different species (*Triturus cristatus*, *Lissotriton helveticus*, *Ichthyosaura alpestris*, Supplemental Table 1). Two skin swabs per individual were collected at three different sites during local monitoring projects between 2020 and 2023 with one set of swabs undergoing kit-based DNA extraction soon after collection (hereafter, “Kit”; Qiagen Blood & Tissue following (27) or PrepMan Ultra following the manufacturer’s instructions) and the other set of swabs being stored at −20C° until “quick” DNA extraction (hereafter, “Quick”; Fig. 1; 21) in Fall 2023. This selection of field-collected samples enabled us to determine FINDeM’s LOD, to assess a quick extraction technique that allows on-site diagnostics, and to account for variation between host species and possibly distinct *Bsal* lineages (28). Additionally, we used sterile swabs and DNA extracts of cultured zoospores as negative and positive controls, respectively. FINDeM reactions were performed as described above while qPCR reactions of the same isolates were performed in duplicate following (29). A full protocol for performing FINDeM in a limited equipment situation can be found in Supplemental File 2.

**Figure 1:**
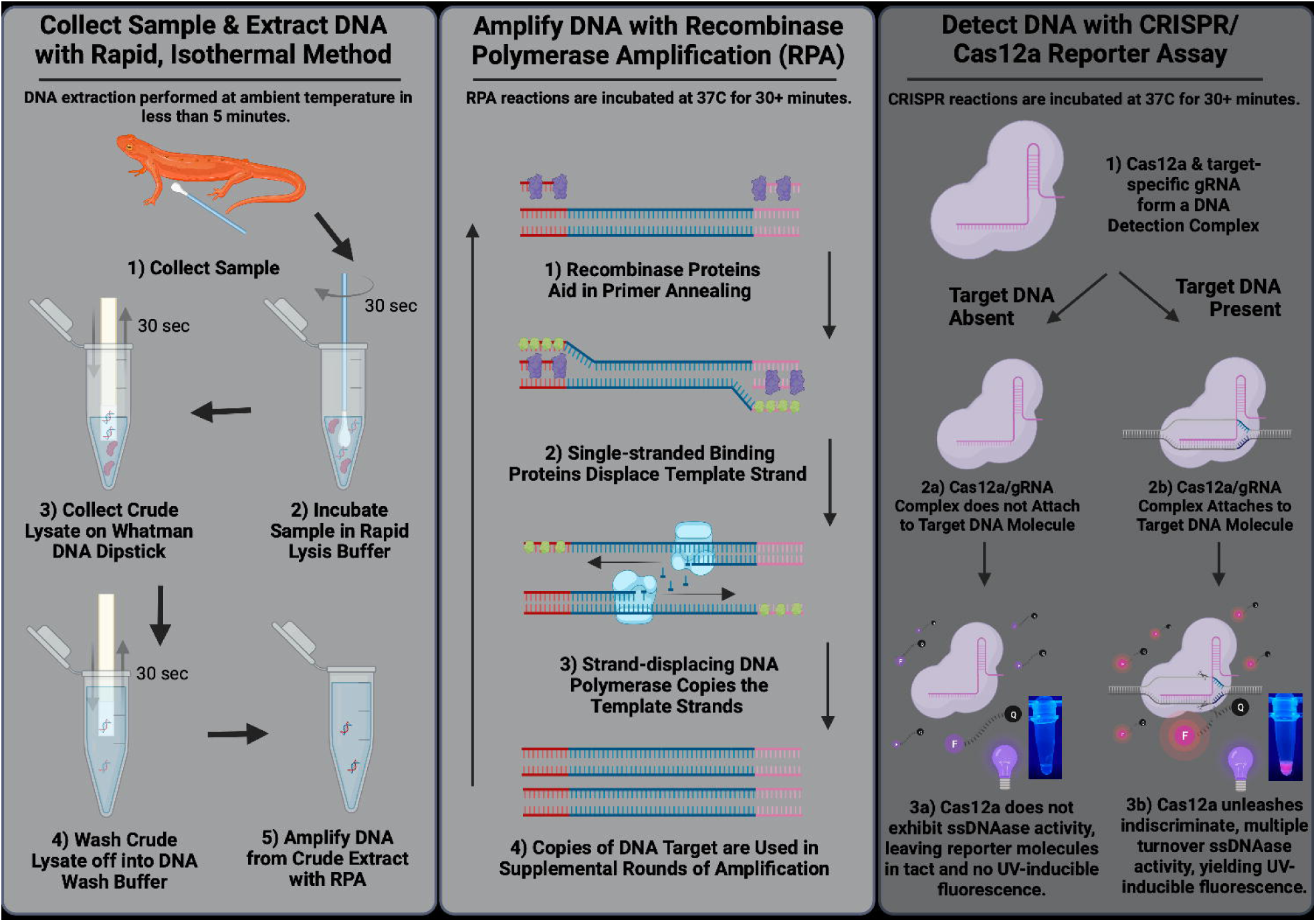
Methodological overview of *Bsal*-targeting FINDeM assay that includes sample collection and DNA extraction (panel 1), DNA amplification with recombinase polymerase amplification (panel 2), and DNA detection using a CRISPR/Cas12a fluorescent reporter assay (panel 3).

### Data processing and analysis

The qPCR-based copy number estimates and ROX-channel relative fluorescence in the *Bsal* DNA standards experiment were derived from the default analysis of the QuantStudio 3 software while copy number estimates of field swab samples were derived from the default analysis of a StepOnePlus software (both Applied Biosystems™; Waltham, MA, USA). FINDeM-based detection statuses for the *Bsal* standards experiment were assessed for unambiguous fluorescence and the threshold for detection was retroactively set to 500,000 relative fluorescence units (RFU) based on these observations. Raw RFU values were then used in two generalized additive models (function: mgcv::gam, formula: RFU ∼ s(log10(Bsal copies + 1), method = ‘REML’; 30), one using data only from ZGE standards and the other only using data from plasmid standards. FINDeM-based *Bsal* detection statuses for swab samples were also assessed for fluorescence by two observers and were separated into four groups: Positive (i.e., bright, unambiguous fluorescence when compared to controls), Slight Positive (i.e., dull, but unambiguous fluorescence when compared to controls), Inconclusive (i.e., dull fluorescence that did not unambiguously differ from that of negative controls), and Negative (i.e., no fluorescence). All images of FINDeM results are available in Supplementary File 1 for external verification.

## Results

### FINDeM validation with Plasmid and Zoospores DNA Standards

Using FINDeM we were able to detect *Bsal* DNA in both replicates of every sample (i.e., ‘total agreement’) containing 100 or more GEs and 2600 or more plasmid copies, while qPCR was able to detect *Bsal* DNA in every replicate with 1 or more ZGEs or 2.6 or more plasmid copies (Fig 2). However, when requiring only one of each standard’s two replicates to be positive (hereafter, ‘partial agreement’), qPCR detected *Bsal* DNA from a sample containing 0.1 ZGEs while FINDeM detected *Bsal* DNA from a sample of 10 ZGEs and another containing 26 plasmid copies (Fig 2). Generalized additive models indicated that there was a significant correlation between qPCR-estimated DNA copies in a sample and the final emitted ROX fluorescence in FINDeM assays for both the ZGE standards (Fig 2; adjusted R^2^ = 0.966, deviance explained = 97.6%, *p* < 0.001) and plasmid standards (adjusted R^2^ = 0.913, deviance explained = 93.7%, *p* < 0.001).

**Figure 2:**
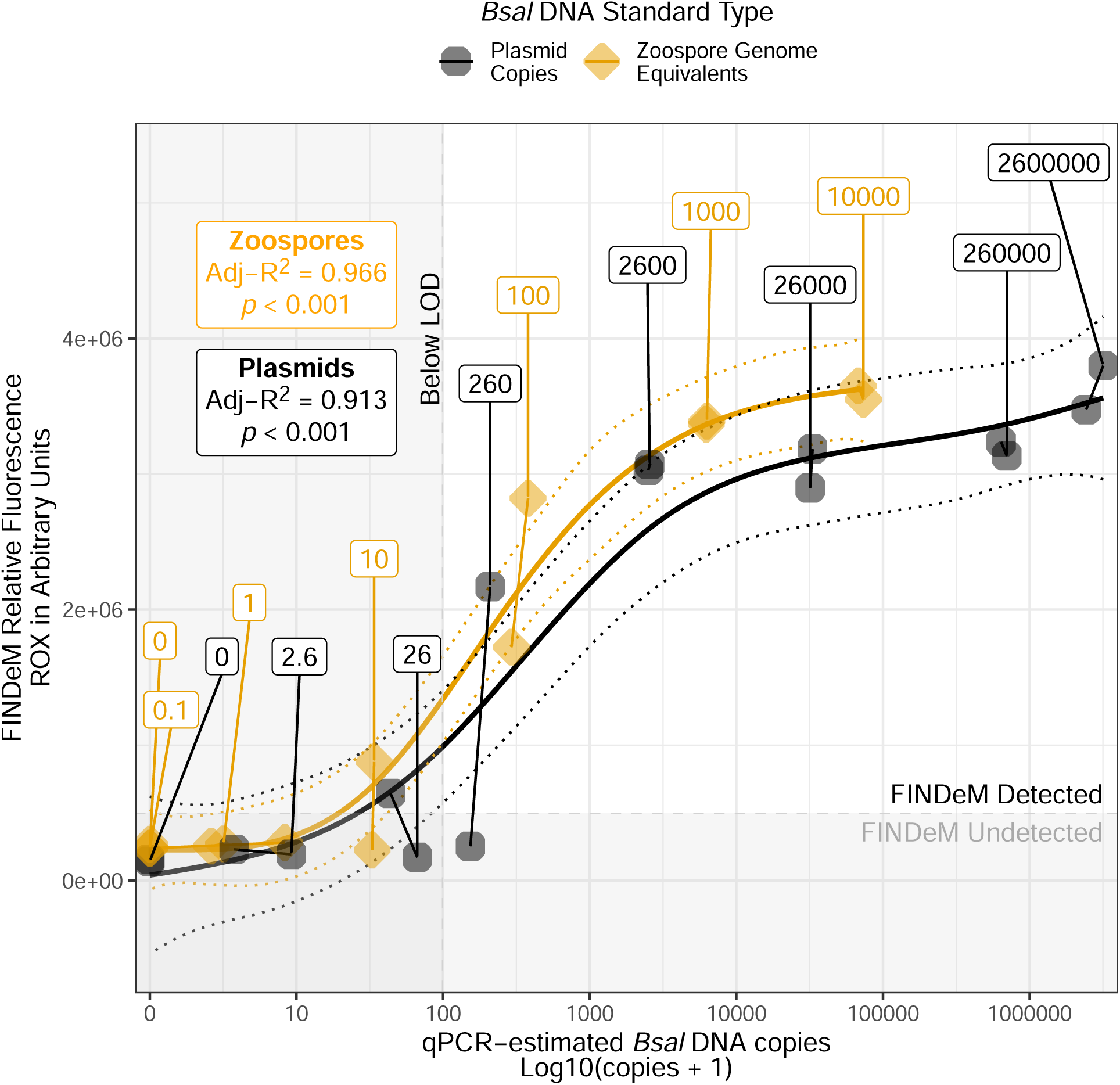
Comparison of Log10, qPCR-estimated *Bsal* DNA copies in plasmid standards (black circles) and DNA extracted from zoospores (orange triangles) to ROX fluorescence emitted by the *Bsal* FINDeM assay. A vertical line at 100 DNA copies indicates the commonly accepted qPCR limit of detection (14) while a horizontal line at 500,000 fluorescence units indicates the threshold of fluorescence for *Bsal* positive samples within this study. Smoothing lines follow the predictions of generalized additive models for plasmid standards (black lines) and zoospore standards (orange line) and the relevant model statistics for each can be found in labels in the top left of the figure.

### FINDeM validation with field samples

We then compared the degree of agreement between FINDeM and qPCR on kit-extracted field swabs and found the results agreed between 70.5% of replicates (Fig 3; n = 44 replicates; 27 agreed positives, 4 agreed negatives, and 13 FINDeM false negatives). When considering the commonly accepted qPCR LOD of 100 copies (14), this resulted in both partial and total agreement of 86.4% between paired samples (n = 22 newts). In the same comparison using quick-extracted swabs, we once again found 70.5% agreement between FINDeM and qPCR (Fig 3; 9 agreed positives, 22 agreed negatives, 7 FINDeM false negatives, and 6 qPCR false negatives) resulting in 100% partial agreement and 68.2% total agreement among samples. This markedly lower degree of total agreement was caused by FINDeM missing one replicate of a qPCR-positive sample (Fig 3; FIN9) as well as FINDeM detecting *Bsal* in 3 positive samples (as verified from kit-extracted corresponding samples) that qPCR did not detect (Fig 3; FIN3, FIN18, & FIN21) and another qPCR-positive sample that had DNA concentrations below the 100-copy LOD (Fig 3; FIN12).

**Figure 3:**
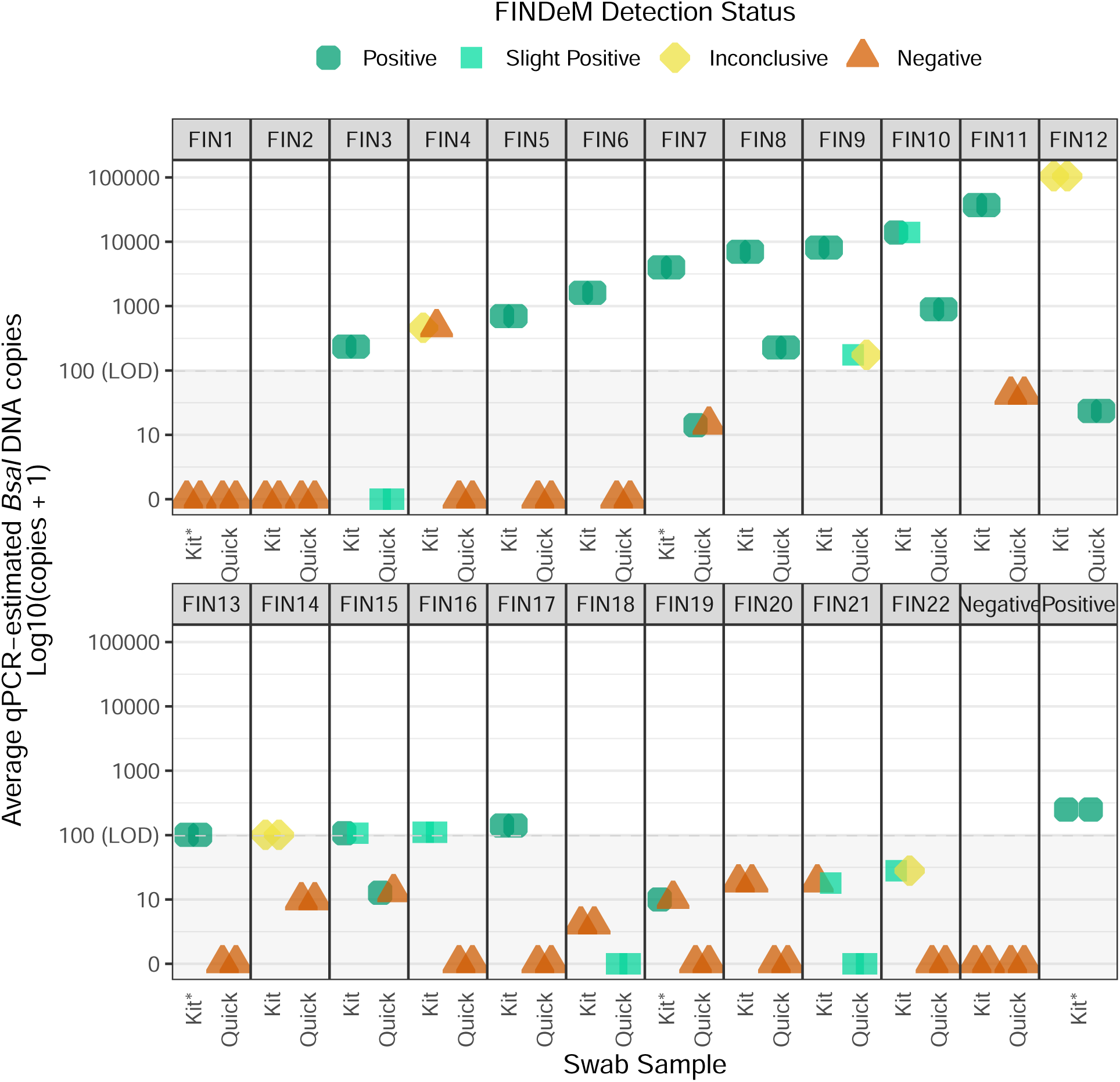
Results of FINDeM on field-collected samples (colored shapes) compared to average *Bsal* DNA estimates from qPCR. The commonly accepted limit of detection is denoted by a dotted horizontal line and grey coloring below 100 *Bsal* copies. Kit-extracted samples FIN1, FIN7, FIN13, and FIN19 (denoted with an asterisk) were repeated alongside the positive control as the initial results from these samples indicated a failed assay. The images of each sample’s assays can be found in Supplemental File 1.

We then compared results stemming from FINDeM on quick-extracted swabs – the most cost-effective and accessible combination in this study – to results from the standard approach of qPCR of kit-extracted samples. In this comparison, we found 43.2% agreement among swabs (Fig 3; 15 agreed positives, 4 agreed negatives, and 25 FINDeM false negatives) which resulted in 54.5% partial agreement and 40.9% total agreement between paired swabs. To determine if this relatively low level of agreement was related to long-term storage or the quick-extraction protocol, we compared qPCR results between swabs extracted with kit-based or quick methods and found 45.5% agreement between quick-extracted and kit-extracted paired replicates (Fig 3; 6 agreed positives, 14 agreed negatives, 24 quick-extracted qPCR false negatives).

## Discussion

### Validation

In this study, we successfully demonstrated the efficacy of FINDeM for *Bsal* detection against traditional qPCR approaches using two types of DNA standard and field-collected swabs. Although detection sensitivities for each approach varied among replicate swab samples, we show that FINDeM can not only detect *Bsal* below the current LOD of 100-copies (14), but may also be able to detect DNA in *Bsal*-positive samples that go undetected by qPCR; possibly owing to Cas12a’s ability to return higher fluorescence per target molecule than the one-to-one ratio of target molecule to fluorescence in qPCR (19). Additionally, based on the observed correlation between ROX fluorescence and estimated DNA copies by qPCR, it also appears that FINDeM may allow for coarse quantification of *Bsal* DNA when properly optimized and used alongside the correct controls.

However, we also uncovered discrepancies between paired swab samples of the same individual that underwent different extraction methods. In agreement with that described in (19), we found that the ‘quick’ DNA extraction approach was less effective than standard, kit-based DNA extractions, leading to substantially lower rates of agreement between qPCR-analyzed, kit-extracted samples and quick-extracted samples with both qPCR (45.5%) and FINDeM assays (43.2%). However, while (19) found that rapid DNA extractions of serially diluted *Bd* ZGE standards were only one order of magnitude less effective than kit-based methods and still allowed for the qPCR-based detection of hundreds of DNA copies, we frequently observed that copy estimates with quick-extracted, *Bsal*-positive samples were multiple orders of magnitude lower than their kit-extracted counterparts or didn’t return qPCR-detectable concentrations of DNA at all. As it appears unlikely that the extractability of DNA would differ greatly between fungal species within the same genus, we believe that this discrepancy may have come as a result of DNA degradation caused by the storage of quick-extracted swabs at −20C°. Nevertheless, while the combination of FINDeM with kit-based *Bsal* extraction methods performed quite well, we suggest that researchers critically consider the extraction method used alongside FINDeM, with those samples having undergone long-term storage (e.g., this study) or being collected from low-template (e.g., eDNA) or low-quality samples (e.g., formalin-fixed museum samples) requiring the greatest scrutiny.

### Application and recommendations

Though our findings support the applicability of FINDeM in future *Bsal* studies, we provide recommendations on how to best obtain results that are comparable to those stemming from established, qPCR-based methods. In *Bsal* qPCR diagnostics, it is standard to run samples in duplicate, concluding that a sample is positive only if both replicates are positive above a certain copy-number threshold (e.g., 100 copies; 31). For FINDeM, we also suggest the analysis of two technical replicates to account for the stochasticity observed in the low volume RPA and CRISPR reporter assays. However, as FINDeM did not return a single false positive result in this study and does not allow for the strict quantitative cutoffs that are possible with qPCR, we recommend that samples returning even a single positive replicate be treated as *Bsal*-positive when using FINDeM. Although this may lead to higher instances of sub-LOD false positives, researchers can still perform qPCR of these samples in light of this LOD and may even decide to only perform qPCR on the samples that were FINDeM positive, thus conserving time and resources by not attempting to quantify presumed *Bsal*-negative samples. Alternatively, researchers may elect to perform qPCR only on FINDeM *Bsal*-negative samples or even batches of such samples, using positives from the more rapid (1hr vs. qPCR ∼2hr), accessible, and cost-effective FINDeM reaction (∼$1.30/reaction vs. qPCR ∼$4.30/reaction) as an initial means of detection and subsequent qPCR only to ensure that no positive samples were missed. We also highlight that the interpretation of FINDeM when performed without a fluorimeter is dependent upon the observer and recommend that images of final reaction tubes be collected and available for later verification. Finally, it is essential that appropriate positive (e.g., zoospore cultures) and negative controls (e.g., sterile swabs) be assessed alongside samples to not only monitor the efficiency of DNA extraction, amplification, and detection reactions, but to also provide frames of reference for the fluorescent signals emitted by FINDeM positive and negative samples.

### Future directions

Limited funding has prevented large-scale screening of *Bsal* in most member states of the European Union, despite the imminent risk of undetected disease outbreaks for Europe’s salamander diversity (32). However, due to its relatively low cost, FINDeM may prove to be an essential supplemental or even alternative method to qPCR, especially when used to trace the expansion and prevalence of *Bsal* – two tasks that likely do not require knowledge of pathogen load, which is one of the main benefits of qPCR. Moreover, the ease of use and minimal technical expertise required for FINDeM increases its potential within citizen science scenarios. While the direct benefits of citizen scientists using FINDeM to increase the depth and breadth of *Bsal* sampling are obvious, there are also indirect benefits of engaging the public within such research: by designing a method that allows the public to be active stewards of the species within their environment, we can also shed greater light on the importance of biodiversity conservation and thus warrant greater resource contributions to support important ecological research.

Although active and passive surveillance help to understand *Bsal* spread, few tools are yet developed for identifying and mitigating *Bsal in situ* in the long-term (16). This is especially important for some newt and anuran species that can carry *Bsal* without expressing clinical signs of infection; essentially serving as *Bsal* reservoirs that can act as ‘silent spreaders’ of the pathogen to more sensitive species (33). FINDeM now allows researchers to identify infected specimens directly in the field with molecular sensitivity, which might be needed for several containment strategies such as drug-based *in situ* treatments (27) or even removal of infected hosts (16). Further, *in situ* detection may also help to selectively collect *Bsal* isolates only from confirmed-infected hosts, thus limiting the effort spent trying to isolate *Bsal* from uninfected individuals while enabling a deeper understanding of the dispersal and evolution of localized *Bsal* isolates which have been shown to vary greatly in their ecological niche (28).

Conservationists have repeatedly called for more “biosecure” amphibian trade for regions that appear *Bsal* naïve, such as in the Americas where the introduction of *Bsal* is expected to cause widespread population declines in the world’s greatest salamander diversity hotspot (15). Although imports of salamanders to the US have since been regulated, known anuran hosts of *Bsal* are not subject to such regulations (34) and the European Commission repealed the trade ban of salamandrids in 2022. Therefore, we recommend that responsible authorities consider the use of FINDeM and other rapid, CRISPR-based detection methods (e.g., SHERLOCK/Cas13a; 35, 36) on traded amphibians to ensure both pathogen-free trade and protection of communities in pathogen-free refuges.

## Conclusion

To mitigate the impacts of *Bsal* on global amphibian populations, it is imperative that the assays used to detect this fungal pathogen be precise, cost-effective, and accessible to those entrusted with monitoring the spread of this fungal pathogen. However, employing the sensitivity of DNA-based pathogen diagnostics has long required dedicated laboratory spaces, expensive equipment, and specialized training in molecular biology, which alongside the considerable lag time to diagnosis has inhibited our ability to make rapid treatment and management decisions for infected individuals. Here, we demonstrate the applicability of FINDeM – a simplified, rapid, and inexpensive method for molecular diagnostics – in the identification of *Bsal* DNA from assay standards and epidermal swabs of wild caudates. By designing a more accessible, yet similarly sensitive, method for *Bsal* detection, it is our hope that a greater number and diversity of individuals will not only be able to contribute to our overall understanding of *Bsal* but serve a key role in its mitigation and restriction of further spread across the globe.

## Acknowledgements

This work was supported by the Frank J. Schwartz Early Career Research Fellowship (Pymatuning Lab of Ecology) awarded to Brandon D. Hoenig and by the RIBBiTR Biological Integration Institute led by Corinne L. Richards-Zawacki (NSF; DBI-2120084). Further, we are grateful to Miranda Kosowsky (University of Pittsburgh), Sabine Naber (Trier University), and Karin Fischer (Trier University) for their support in the lab. Permits for field samples were granted by the Struktur- und Genehmigungsdirektion Nord Rheinland-Pfalz (permit number: 425-104-778-0007/2022) and the Eifel National Park. Field sampling was kindly financed by the Ministry of Environment Rhineland-Palatinate within the funding program “Aktion Grün” (102-88 712/2020-37#15).

## Data Availability Statement

The raw data and R scripts required to reproduce our analysis can be found on osf.io upon acceptance and further questions about the data and analysis can be directed to the corresponding author.

## Corresponding author biography

Brandon D. Hoenig is a postdoctoral researcher with the University of Pittsburgh and the RIBBiTR Biology Integration Institute. His research interests are focused on using molecular tools to understand species interactions as well as developing more accessible scientific techniques to increase the diversity of individuals within the STEAM fields.

## Protocol for the field-based detection and quantification of *Batrachochytrium salamandrivorans* DNA from swab samples

Created 23 December 2023

Contact Brandon Hoenig with questions.

### Materials

Rayon Swabs

0.2mL PCR tubes (individual and with lids)

0.2mL PCR tubes (in groups of 5+ and with lids)

<u>For Detection:</u> Prelabel tubes with 1 negative control, 1 positive control, and 3+ samples.

Add 1uL of *Bsal* DNA standard (10,000 zoospore genome equivalents/1uL) & 4uL molecular grade water to positive control.
Add 5uL of molecular grade water to negative control.
Spin down this mixture with centrifuge or salad spinner

<u>For Quantification:</u> Prelabel tubes with 1 negative control, 1 “medium concentration” positive control, 1 “high concentration” positive control and 2+ samples. 0.2mL PCR tubes (in groups of 3 and with lids)

Fill as many groups of 3 as needed with 100uL of Extraction Buffer
Fill as many groups of 3 as needed with 200uL of Wash Buffer.

0.5uL – 10uL single-channel pipette (or a combination of pipettes that cover this range)

0.5uL – 10uL multi-channel pipette (or a combination of pipettes that cover this range)

20uL – 200uL single-channel pipette

Filter tips for 0.5uL – 10uL pipette

Filter tips for 20uL – 200uL pipette

Pipetting can be replaced by using scaled microliter capillaries here.

Single-use laboratory gloves

Salad Spinner or microcentrifuge

96-well PCR rack

Whatman #1 paper cut into hole-punch size (∼3mm diameter) or Whatman #1 strips that are waxed on one end (e.g., using candlewax) to serve as a handle and left unwaxed on the other end (∼3×3mm) to absorb DNA solutions.

Small black sheet of paper as a background

Dark room or homemade dark box

UV Light

Orange UV Glasses

### Reagents

**TwistDx Basic RPA Kit** <u>ordered from TwistDx</u>

Lyophilized RPA Master Mix

Rehydration Buffer

280mM Magnesium Acetate (MgOAC)

10uM SterF primers – 5’ TGCTCCATCTCCCCCTCTTCA 3’

10uM SterR primers – 5’ TGAACGCACATTGCACTCTAC 3’

**Extraction Buffer; 20mM Tris HCl pH 8.0, 25mM NaCl, 2.5mM EDTA, 0.05%SDS, 2% PVP-40**

For 50mL:

1mL 1M Tris

250uL 5M NaCl

250uL 0.5M EDTA

0.025g SDS (ours were stringy SDS, not pellets)

1g PVP-40

Fill to 50mL with nuclease-free water

**Wash Buffer; 10mM Tris HCl pH 8.0**

**1uM LbCas12a** Ordered from NEB

**1.25uM Bsal Guide RNA** - 5’ rUrArArUrUrUrCrUrArCrUrArArGrUrGrUrArGrArUrArGrArArUrCrArArUrArGrUrArUrUrUrUrCrUr UrG 3’ Ordered from IDT (See help section on this page if adapting for other uses).

**25uM 5’-ROX-NNNNNNNNNNNN-BHQ2-3’ ssDNA** <u>Ordered from IDT</u>

### Prior to sampling

**RPA Reaction Buffer (5, 4.5uL reactions per tube – *add to positive control tube last*):**

Add 17.7uL of Rehydration buffer to a TwistAmp Basic Tube (white powder)

Add 4.8uL of 20uM ITS/5.8S primer mix (2.4uL of each) to tube

Vortex/pipette up and down vigorously until lyophilized powder is in solution. <u>This mixture will be viscous.</u> Spin down in centrifuge (or salad spinner).

Add 4.5uL of the RPA Mix (22.5uL in total) to each of the 5+, connected 0.2mL PCR tubes. Be certain to add to pre-labelled negative control tube first, the 3+ sample tubes next, and the positive control tube last. Leave tubes at room temperature until sampling event later that day. If longer term storage is needed, then freeze and centrifuge before use.

**CRISPR Reaction Buffer (per reaction; add on 1 extra reaction per 5 reactions)**

**<u>For 1 reaction</u>**

Add 1uL of NEBuffer 2.1

Add 0.5uL of 25uM ROX-12N-BHQ2

Add 0.5uL of 1.25uM *Bsal* gRNA

Add 0.5uL of 1uM LbCas12a

Spin down in centrifuge (or salad spinner) and allow CRISPR Reaction Buffer to incubate for 20 - 30 minutes at room temperature. During this step, the gRNA and LbCas12a are forming a DNA detection complex.

After 20 - 30 minutes, aliquot 2.5uL of CRISPR Reaction Buffer to each of 5 pre-labelled and connected 0.2mL PCR tubes.

Store at room temperature until sampling later that night. If storing long-term, store in freezer and spin down prior to use.

**Extraction Buffer**

Add 100uL of extraction buffer to each of a set of 3 connected PCR tubes.

**Wash Buffer**

Add 200uL of wash buffer to each of a set of 3 connected PCR tubes.

**Whatman #1 Filter Discs (3mm) or Whatman #1 ‘DNA Dipsticks’**

Use a standard hole-punch to punch out 3mm Whatman #1 filter discs or Fold Whatman #1 paper in half, dip all but 3mm of the non-folded side into candlewax, and slice into 3mm wide strips. Most of the dipstick will be waxed and will serve as a handle, but a 3×3mm unwaxed area will be used as a replacement for the Whatman #1 Filter Disc.

### When in the field

#### *Bsal* DNA Detection Protocol

Using fresh gloves, open the Rayon swab and swab the amphibian 40 times (5 times on back, 5 times on belly, 5 times on each hand, 5 times on each foot, 5 times on each hindlimb groin area).

Place swab into 100uL of Extraction Buffer and agitate swab up-and-down and across the inside of the tube vigorously for 30 seconds. Be careful not to spill between sample tubes.

Place used swab into labeled 1.5mL or 2mL tube for long-term storage.

Place Whatman #1 disc or unwaxed side of DNA dipstick into DNA extraction buffer. Agitate the Whatman #1 circle/dipstick in the buffer vigorously for 30 seconds (a pipette tip works well for the circles).

Using a pipette tip and a fresh glove transfer the Whatman #1 disc or unwaxed side of DNA dipstick to 200uL of wash buffer and agitate for 30 seconds. Be careful not to spill between sample tubes.

Using the single-channel pipette, add 5uL of wash buffer (i.e., DNA extract) to each of the three ‘Sample’ RPA reaction tubes. Positive control DNA (∼10000 copies/5uL) and negative control (nuclease-free water) should have been added ahead of time in a largely DNA free space.

Using the multi-channel pipette (or a single channel pipette on the wall of the PCR tube), add 4.5uL of RPA Reaction Mix to the ‘Sample/Standard’ PCR tubes. You may decide to add the 0.5uL of 280mM MgOAc to each RPA reaction tube before the RPA mix in the lab or, you can add 0.5uL of 280mM MgOAc to the side wall of the reaction after 4.5uL of RPA Reaction Mix has been added.

Using a centrifuge or salad spinner, ensure that all reaction liquid makes it to the bottom of the tube at the same time. Alternatively, you can use a pipette or pipette tip to ensure that all reaction volume is in the bottom of the tube. *<u>However, It is critical that MgOAc is added to each tube at the same time as the reaction initiates the moment that MgOAc is added to the RPA Reaction Mix</u>*.

Incubate tube at 37C (taping in armpit and closing arm by side is easiest in my experience) for 30 minutes. Note: Handheld incubation does not get tubes warm enough for rapid amplification reaction.

After incubation and using a multi-channel pipette, add 2.5uL of the CRISPR Detection Mix to the 10uL of completed RPA Reaction. **<u>It is advised against to pipette the RPA product into the CRISPR Detection Mix as the amplified DNA from RPA may contaminate subsequent reactions if it makes its way into the pipette. In our experience, filter tips may not help to reduce this contamination possibility.</u>** The 2.5uL of CRISPR Detection Reaction can be added to the side of each tube and spun down in a salad spinner/centrifuge to ensure that the 2.5uL contacts the RPA reaction at the same time.

Incubate tube at 37C (taping in armpit and closing arm by side is easiest in my experience) for at least 15 minutes. Note: Handheld incubation does not get tubes warm enough for rapid detection reaction.

In the dark, visualize with UV light (and UV glasses to reduce glare) against a black background and compare to positive and negative controls. Positives will fluoresce under UV light while negatives will not. If unclear, incubate in arm pit for 5-10 additional minutes and visualize again.

### *Bsal* DNA Quantification Protocol

Follow the same steps as in the Bsal DNA Detection Protocol, however, analyze 2+ amphibian swabs alongside 3+ standards (e.g., 0, 100 zoospore genome equivalents, and 10,000 zoospore genome equivalents; though whatever standards are decided upon should be validated in the lab first) and compare amphibian swab sample results to standards for coarse quantification.

